# A spontaneously blinking fluorescent protein for simple single laser super-resolution live cell imaging

**DOI:** 10.1101/200014

**Authors:** Yoshiyuki Arai, Hiroki Takauchi, Yuhei Ogami, Satsuki Fujiwara, Masahiro Nakano, Takeharu Nagai

## Abstract

Super-resolution imaging techniques based on single molecule localization microscopy (SMLM) broke the diffraction limit of optical microscopy in living samples with the aid of photoswitchable fluorescent probes and intricate microscopy systems. Here, we developed a fluorescent protein, SPOON, which can be switched-off by excitation light illumination and switched-on by thermally-induced dehydration resulting in an apparent spontaneous blinking behavior. This unique property of SPOON provides a simple SMLM-based super-resolution imaging platform which requires only a single 488 nm laser.

## Main text

### Introduction

Super-resolution imaging techniques are now indispensable to explore biological structure at nano-meter scales in living cells^1^. To break the diffraction limit many super-resolution methods have been developed. These include stimulated emission depletion (STED)^2^, reversible saturable optical fluorescence transition (RESOLFT)^3^, photoactivation localization microscopy (PALM)^4^, stochastic optical reconstruction microscopy (STORM)^5^, and structured illumination microscopy (SIM)^6^. Initially, only PALM and STORM required photoswitchable fluorescent probes. But now, many other super-resolution techniques use photoswitchable fluorescent probes to reduce illumination power density and improve biocompatibility^7^. Genetically encoded photoswitchable fluorescent proteins (PSFPs) are especially promising as they can easily be targeted to the proteins of interest in living cells^8^. Among PSFPs, Dreiklang was the first reported PSFP which can be repeatedly switched on and off by using three different wavelengths of light (360 nm, 405 nm, and 515 nm or 488 nm)^9^. In addition, Dreiklang can switch on spontaneously by thermal relaxation, which enables a simpler single molecule localization based microscopy, called DSSM (decoupled stochastic switching microscopy)^10^. In DSSM, the on-state of Dreiklang is thought to transfer to a long-lived dark state by 488 nm irradiation during observation, which makes this strategy similar to ground state depletion microscopy followed by individual molecule return (GSDIM)^11,12^. Compared to GSDIM, DSSM enables super-resolution imaging at relatively lower power density illumination for switching off (0.2 kW/cm^2^ by 405 nm light, while GSDIM required 3 kW/cm^2^). However, DSSM still requires 405 nm illumination to make Dreiklang adopt the fluorescent off state. Recent studies have highlighted the phototoxicity of 405 nm irradiation during super-resolution imaging which hampers biocompatiblity^13^. In addition, the requirement for close coordination of two laser lines for super-resolution imaging increases system cost, and complexity, which may be barriers to adoption among non-experts.

To overcome this problem, we developed an improved PSFP that has a fast blinking behavior due to thermal switching ON, and rapid photo switching OFF with the same 488nm excitation light needed for fluorescent emission. This enables simple super-resolution imaging using single wavelength laser illumination.

## Results

### Development of SPOON

To develop a fast photoswitching mutant of Dreiklang, we performed extensive random mutagenesis using the Dreiklang gene as a template by error-prone PCR and DNA shuffling ^14^. We transformed bacteria with the mutant cDNA library and screened ∼50,000 colonies for fast switching mutants of Dreiklang. Ultimately, we obtained a mutant carrying five substitutions: I47V, T59S, M153T, S208G, and M233T (**Supplementary Figure 1**). This mutant exhibited an absorption spectrum almost identical to Dreiklang during on and off states (**Supplementary Figure 2**). The absorption spectrum in the thermal equilibrium state (switched on) showed two peaks at 411 nm and 510 nm, and an emission peak at 527 nm. (Figure 1a, Table 1). The absorption peaks are responsible for photoswitching off and excitation, respectively. Upon photoswitching to the off state by 438 nm irradiation, a new absorption peak at around 339 nm appeared which is responsible for photoswitching on (Figure 1b, **Supplementary Figure 3**). Fluorescence quantum yield (QY) of the mutant and Dreiklang were 0.50 and 0.44, respectively, with a molar extinction coefficient at 510 nm of 54,000 M^-1^cm^-1^ and 81,000 M^-1^cm^-1^ in the mutant and Dreiklang respectively (Table 1), suggesting that the brightness of mutant was slightly lower than Dreiklang itself. We named this Dreiklang mutant as SPOON, *<u>SPO</u>*ntaneous switching *<u>ON</u>* fluorescent protein.

**Table 1.** Characteristics of SPOON and Dreiklang

| Parameters | SPOON | Dreiklang |
| --- | --- | --- |
| Wavelength of switching off light (nm) | 339 | 339 |
| Wavelength of switching on light (nm) | 411 | 412 |
| Fluorescence excitation peak (nm) | 510 | 510 |
| Fluorescence emission peak (nm) | 527 | 527 |
| Fluorescence quantum yield | 0.50 | 0.44 |
| Extinction coefficient@510 nm ($M^{-1} \text{ cm}^{-1}$) | 54,000 | 81,000 |
| Extinction coefficient@405 nm ($M^{-1} \text{ cm}^{-1}$) | 31,000 | 19,000 |
| Extinction coefficient@360 nm ($M^{-1} \text{ cm}^{-1}$) | 24,000 | 22,000 |
| Time constant of photoswitching off (s) | 0.77 (83.3%)<br>3.43(16.6%) | 3.97 |
| Time constant of photoswitching on (s) | 4.84 | 5.93 |
| Time constant of spontaneous switching on (s) | 144.9 | 332.8 |
| Time constant of maturation speed (hours) | 1.86±0.15 | 2.63±0.09 |

**Figure 1.**
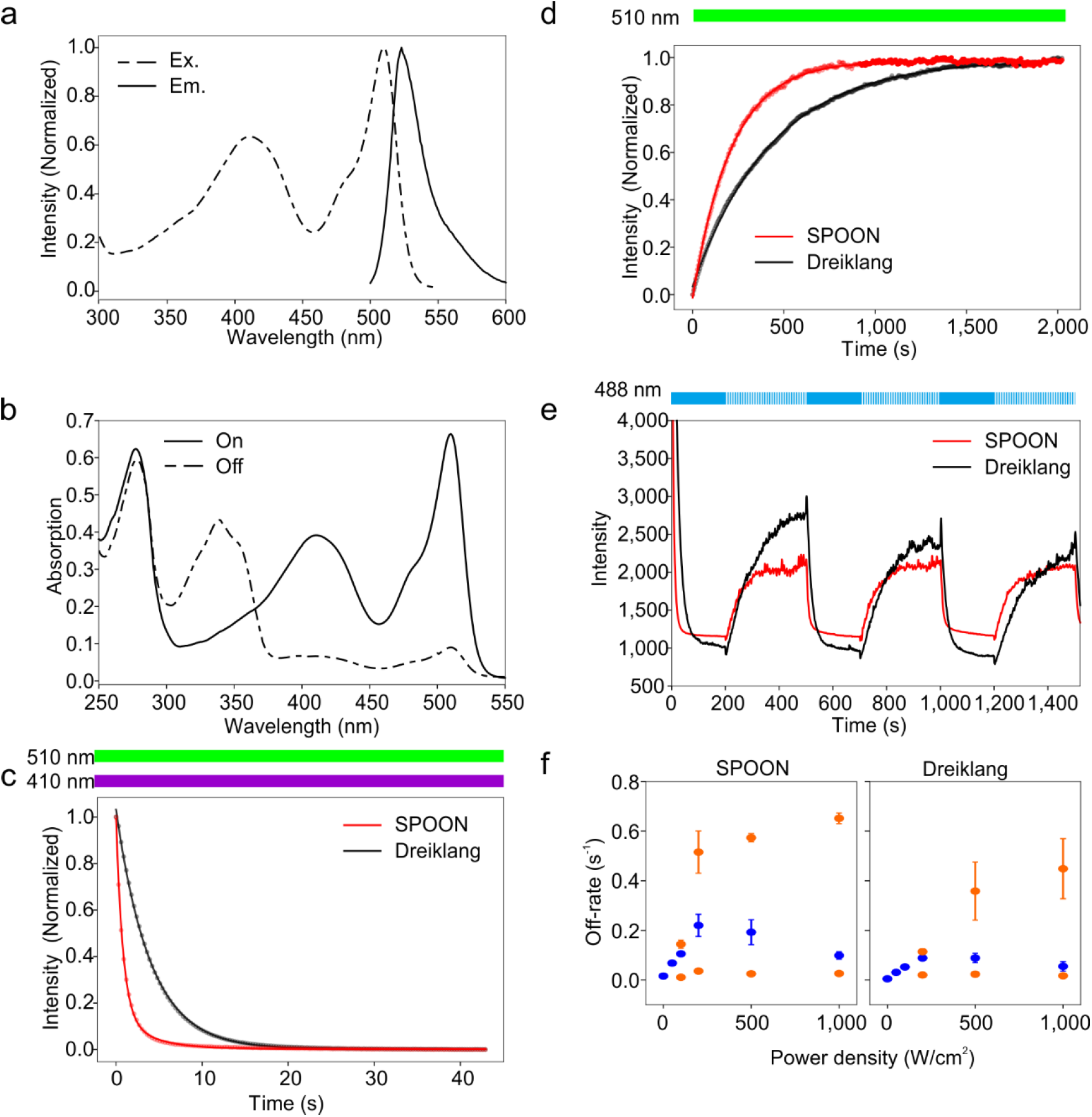
Comparison of characteristics between SPOON and Dreiklang. a, Absorption and emission spectra of SPOON. **b**, Absorption spectra of SPOON in on and off states. **c,** Photoswitching-off kinetics under continuous 488nm and 510nm irradiation) of SPOON (red) and Dreiklang (black). For Dreiklang, data were fitted by single exponential decay curve, while SPOON data was fitted by a double exponential decay curve. **d,** Thermalswitching-on kinetics of SPOON and Dreiklang under continuous 510nm irradiation **e.** Photoswitching-off by 488 nm irradiation at 100 W/cm^2^. Timing of 488 nm of continuous (solid line) and periodic (dashed line) are indicated as bar at the above of graph. **f.** Rate constants of switching off. At higher power density, data were fitted by double exponential curve (orange) instead of a single one (blue). Error bars indicates standard deviation (n=2∼4).

**Figure 2.**
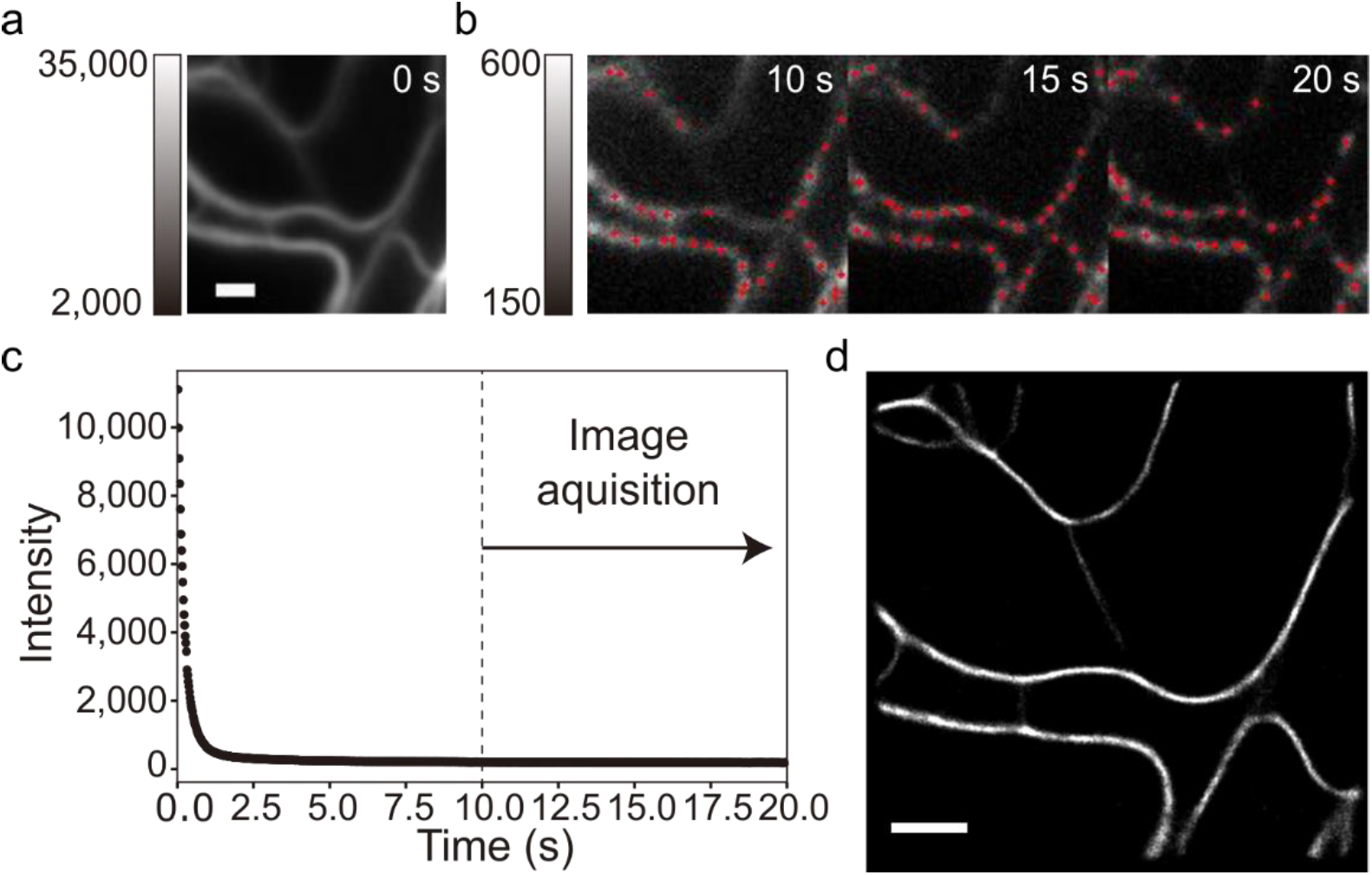
SMLM with only 488 nm irradiation. **a.** The first frame image of Vimentin-SPOON **b.** Montage images taken every 100 frames after 10 s. Red dots indicated the detected molecules by ThunderSTORM software. **c.** Time trajectory of average intensity. Arrow indicates the duration for super-resolution image reconstruction shown in **d**. **d.** Reconstructed super-resolution image using images obtained after 10 s irradiation. Scale bars is 1 μm.

### Characteristics of SPOON

The photoswitching kinetics of SPOON using purified protein embedded in a 15% polyacrylamide gel was examined. Fluorescence signals during photoswitching off were recorded upon 410 nm irradiation in the presence of excitation light (510 nm). The time constant of photoswitching off for SPOON was five times faster than that of Dreiklang (SPOON; 0.77 s vs Dreiklang; 3.97 s) (Figure 1c and Table 1). The speed of photoswitching on in SPOON was slightly faster than that of Dreiklang (4.84 s and 5.93 s, respectively) (**Supplementary Figure 4**). The time constant of thermal switching on for SPOON was approximately 2.5 times faster than that of Dreiklang (∼144.9 s and ∼332.8 s, respectively) (Figure 1d). Photoswitching fatigue resistance of SPOON was 1.7 times longer than that of Dreiklang (**Supplementary Figure 5**). Next, we tested the oligomeric states, maturation time, and pH stability of SPOON. Both SPOON and Dreiklang showed a dimeric tendency in gel filtration chromatography (**Supplementary Figure 6**), whereas semi-native polyacrylamide gel electrophoresis showed monomerization (**Supplementary Figure 7**) consistent with prior reports^9^. The chromophore of SPOON showed an almost identical p*K*a value to Dreiklang (**Supplementary Figure 8**), with a maturation time constant of 1.86±0.15 hours (n=3, mean±s.e.m.) at 37°C (**Supplementary Figure 9**), which was 1.4 times faster than that of Dreiklang (2.63±0.09 hours, n=3, mean±s.e.m.). Although the brightness of SPOON is slightly lower than that of Dreiklang according to its molar extinction coefficient and fluorescence quantum yield (Table 1), photon numbers and localization uncertainty were almost identical to Dreiklang (**Supplementary Figure 10**).

### Photoswitching upon 488 nm excitation

Interestingly, we found that continuous illumination by 488 nm light stimulated photoswitching-off in both Dreiklang and SPOON (Figure 1e). By periodic illumination of 488 nm (100 ms irradiation followed by a 900 ms observation interval), thermal switching-on was seen. These photoswitching-off and thermal switching-on events were observed repeatedly. Photoswitching-off speed depended on the illumination power density. Rate constants of photoswitching-off were obtained by exponential decay curve fitting. At higher power density, time constant of photoswitching-off appeared to be a double, rather than a single, exponential curve, suggesting the existing of two states (Figure 1f). We also evaluated the photoswitching-off and thermal switching-on process by conventional fluorescence microscopy in living cells expressing Dreiklang or SPOON (**Supplementary Figure 10**). 472 nm illumination (15 mW/cm^2^) induced photoswitching-off, whereas 500 nm irradiation allowed the proteins to transit fluorescent-on state. These results suggest that a novel photoswitching-off pathway is present in Dreiklang and SPOON that may not require the transition from an excited state to a long-lived triplet state described for GSDIM since the illumination power density was much smaller than that reported^11^.

### Localization of SPOON in living cells

To test the performance of SPOON in living cells, we fused it with various proteins of interest and expressed these in mammalian cells. Appropriate localization of C terminal-(α-actin, Golgi body-localizing signal, fibrillarin, Cox-VIII, histone H2B, clathrin, and Nup133) and N terminal-fused (paxillin, zyxin, vimentin, β-tubulin, tyrosine-protein kinase, endoplasmic reticulum, and second-mitochondrial activator of caspase) SPOON showed the anticipated expression patterns (**Supplementary Figure 11**). Although both Dreiklang and SPOON were observed as weak dimers in solution by HPLC (**Supplementary Figure 6**), these results suggest that in mammalian cells this property does not cause significant impediment to the normal function of the fusion partner.

### Super-resolution imaging by single wavelength laser illumination

We next investigated if the faster thermal switching-on property of SPOON (Figure 1d), together with the 488 nm triggered photoswitching-off (Figure 1e) and the possibility to excite sub-maximally at 488 nm was sufficient to allow super-resolution imaging based on a single laser. First, we identified a cell expressing Vimentin-SPOON (Figure 2a). The cell was continuously irradiated by 488 nm light to induce photoswitching-off, and induce fluorescence emission. Under these conditions, the fluorescence of SPOON decreased immediately (Figure 2b). After 10 s, single molecules of Vimentin-SPOON became sparse enough to detect with center localization. The signal from single molecules appeared randomly, sparsely and stably during observation (Figure 2c). A super-resolution image was reconstructed from 2,000 frames (Figure 2d).

## Discussion

### Performance of SPOON

Here we report a novel fluorescent protein, SPOON, which exhibits faster thermalswitching-on and photoswitching-off kinetics than that of the parental protein Dreiklang. The photoswitching-off speed of SPOON was five times faster than that of Dreiklang. In the on-state absorption spectrum of SPOON, we found a large peak at around 405 nm (Table 1, **Supplementary Figure 2**) which may contribute to the accelerated photoswitching-off speed.

### Simple biocompatible super-resolution

Although the excitation peak wavelength of both Dreiklang and SPOON is around 510 nm, these proteins can be excited by 488 nm irradiation. We have found that not only 405 nm irradiation, but also 488 nm irradiation leads to fluorescent off states. Spontaneous thermalswitching-on also occurs during illumination. This ‘tug-of-war’ relationship enables many long-term single molecule observations. At higher illumination power density (200 W/cm^2^ or larger), the photoswitching off decay curve was fitted by a double exponential function, indicating the existence of two states (Figure 1f). One of the components in the photoswitching-off rate is related to the transition from the on to the off state. Another may be related to photobleaching. Therefore, upon 488 nm irradiation, three-state transition models could be considered,

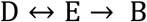

where D is the dark state, E is an emissive state, and B is a bleached state. As a result, SPOON emits photons in blinking manner apparently with only 488 nm illumination. HMSiR, a tetramethyrhodamine derivative, also shows blinking under single wavelength excitation^15^, but as a dye it needs to be conjugated to its target protein outside a cell, or organ which may limit biological application.. However, these kinds of spontaneously blinking fluorophores simplify super-resolution imaging compared to other SMLM based super-resolution techniques (**Supplementary Figure 13**).

In DSSM, or GSDIM, Dreiklang should be irradiated by 405 nm (200 W/cm^2^) or intense 488 nm light (3 kW/cm^2^) to make the on state switch off. Such strong UV illumination damages cells^13^. In this work, because the switching off feature of Dreiklang and SPOON could be triggered by 488 nm irradiation, SMLM could be performed with only one laser (Figure 2). This improves bio-compatibility of super-resolution imaging. Moreover, since SPOON is a decoupled type of PSFP, SPOONs advantages should also be apparentin other super-resolution techniques such as RESOLFT, PALM, and super-resolution optical fluctuation imaging (SOFI).

### Accession code

The SPOON nucleotide sequence has been deposited in GenBank under the accession code LC320161.

## Acknowledgements

We thank Matthew J. Daniels for his critical reading of this manuscript. This work was supported by a Grant-in-aid for Scientific Research on Innovative Areas, ‘Spying minority in biological phenomena (No. 3306)’, from the Ministry of Education, Culture, Sports, Science and Technology, Japan (MEXT) (No. 23115003) (T.N.) and CREST (T.N.).

## Author contributions

T.N. conceived the project. H.T., Y. A., Y. O., and S. F. constructed and characterized SPOON, Y.A. set up the microscopy, H.T. and Y.A. performed imaging. T.N. Y.A., M.N. and H.T. discussed and commented on the results, and wrote the manuscript.

## ONLINE METHODS

### Mutagenesis and cloning

For random mutagenesis, error-prone PCR was performed as follows; 0.5 μL of pRSETB-Dreiklang, 1.5 μL of 10 μM forward and reverse primer for Dreiklang, 5 μL of dNTP mixture (2.5 mM of dATP, 2.5 mM of dGTP, 9 mM of dCTP, and 9 mM of dTTP), 2.5 μL of 8 mM of MnCl2, 0.5 ∼ 2 μL of MgSO4, 5 μL of PCR buffer (Takara Bio Inc., Japan), and ddH2O was mixed to 49.5 μL volume. Then, 0.5 μL of rTaq (R001A, Takara Bio Inc.) was added. Subsequently, PCR was performed to amplify and introduce the mutations. To increase the size of the mutation library, staggered extension process (StEP) was performed, following the published protocol^14^. For non-mutagenic PCR reactions, KOD-Plus (Toyobo CO., LTD., Japan) DNA polymerase was used. The products of PCR-amplified and restriction enzyme-digested DNA were purified by agarose gel electrophoresis, followed by DNA isolation using the QIAEX II gel extraction kit (Qiagen, USA). The amplified DNA was digested with restriction enzyme (RE) endonuclease purchased from New England BioLabs or Takara Bio Inc. Ligation was performed by using T4 DNA ligase with 2×Rapid Ligation Buffer (Promega, USA). cDNA sequencing was performed by dye-terminator cycle sequencing using the BigDye Terminator v3.1 Cycle Sequencing Kit (Life Technologies, USA) and an sequencer (Applied Biosystems, USA). Plasmids were purified by standard alkali methods as described elsewhere. All DNA oligonucleotides used for this study were purchased from Hokkaido System Science and all products were used by following the provider’s recommended protocols. All other chemicals were research grade. Oligonucleotides used in this study are listed in **Supplementary Table 1.**

### Screening of fast photoswitching SPOON

PCR products obtained by error-prone and StEP were ligated into the bacterial expression vector pRSETB (Invitrogen) using *BamH*I and *EcoR*I restriction enzyme sites. The ligation products were transformed to *E. coli* (JM109 (DE3)) and homogenously spread on 95-mm agar plates using sterilized glass beads. *E. coli* were incubated for 16 h at 37°C up to optimal emission of fluorescence from each colony. The Dreiklang mutants expressed in bacterial colonies were switched off with 438/24 nm LED light and switched on with Mercury (Hg) lamps using band-pass emission filter (360/30 nm, Omega Optical. Inc, USA) under simultaneous excitation with 512/25 nm LED light in a home-made uniform illumination system^16^. The emission signals passed through a 535/26 nm band-pass filter (Semrock). The time-course of fluorescence intensity was recorded until it reached a plateau with ∼50- to 200-ms camera exposure time. The image sequence was analyzed using ImageJ software. From each plate, 4 or 5 colonies showing faster switching behavior than Dreiklang were picked. For the second screening step, we used an inverted microscopy (TE2000E, Nikon) with a 20×, 0.5 numerical aperture (NA) objective lens (Nikon), and an LED light source (Spectra X Light Engine, Lumencor), and 100W Hg lamps (Mercury Lamphouse Illuminator, Nikon). For excitation, cyan LED light(488/25 nm) was used. Photo switching on and off was achieved Hg lamp illumination passing through excitation filters (360/20 nm (Omega Optical. Inc) and 410/10 nm (Omega Optical. Inc), respectively) under epi-illumination configuration with a dichroic mirror (FF520-Di02-25x36, Semrock). Emission signals were filtered with an emission filter (FF02-531/22-25, Semrock). Fluorescence images were captured by a scientific CMOS camera (ORCA-Flash4.0, Hamamatsu photonics) at 300 ms exposure time. This screening procedure was repeated ten times so as to obtain the final fast-switching mutant, i.e. SPOON.

### Constructs for mammalian cell expression

SPOON cDNA was amplified by KOD PCR and sub-cloned into pcDNA3 (Invitrogen) using *BamH*I and *EcoR*I RE sites. In order to target SPOON to specific sites such as the mitochondria, Golgi body, nucleus, nucleolus, endoplasmic reticulum (ER), and plasma membrane, we replaced Kohinoor^16^ with SPOON in pcDNA3-CoxVIIIx2–Kohinoor(mitochondria), pcDNA3-GT-Kohinoor (Golgi body), pcDNA3-Kohinoor-H2B (histone 2B), pcDNA3-Kohinoor-Fibrillarin (nucleolar protein), pcDNA3-Vimentin-Kohinoor, and pcDNA3-Zyxin-Kohinoor using *BamH*I and *EcoR*I RE sites^16^, in pcDNA3- Kohinoor-Actin using *Hind*III and *Kpn*I RE sites, in pcDNA3-clathrin-Kohinoor using *Nhe*I and *Bgl*II, pcDNA3-ER-eNanolantern (endoplasmic reticulum) using *BamH*I and *Kpn*I RE sites^17^, and pcDNA3-Lyn-SuperNova using *BamH*I and *Xho*I RE sites^18^. For Nup133 localization, we replaced the β-actin sequence with the Nup133 sequence in pcDNA3-SPOON-β-Actin vector using *BamH*I and *EcoR*I RE sites. For second mitochondria-derived activator of caspase (Smac), we replaced the vimentin sequence with the Smac sequence in pcDNA3-vimentin-SPOON vector using *Hind*III and *Kpn*I RE sites. For β-Tubulin localization, we replaced the Kohinoor moiety of pcDNA3-β-tublin-Kohinoor with the SPOON sequence using *EcoR*I and *Not*I RE sites, 23 amino acid linker peptides were inserted between the β-tubulin and SPOON sequences.

### Protein expression and purification

The SPOON coding sequence was sub-cloned into the pRSET_B_ bacterial expression vector using *BamH*I and *EcoR*I RE sites. SPOON fused with polyhistidine tag was expressed in *E.coli* (JM109 (DE3)) at 23°C for 60 hours in LB bacterial growth medium supplemented with 0.1 mg/ml carbenicillin. The cultured cells were collected and disrupted with a French press (Thermo Fisher Scientific). The recombinant proteins were purified from the supernatant by using Ni-NTA agarose affinity columns (Qiagen) eluted by imidazone and followed by buffer-exchange (20 mM HEPES, 150 mM NaCl pH7.5) gel filtration (PD-10 column, GE Healthcare).

### Protein characterization

Purified proteins of Dreiklang and SPOON in (10 μM in 20 mM HEPES, 150 mM NaCl buffer, pH 7.4) were switched off using 438/24 nm LED light and switched on with Hg lamps using band-pass emission filter (360/30 nm, Omega Optical. Inc). The absorption spectrum of the purified proteins in the on and off state was measured with a spectrophotometer (V-630BIO, JASCO). The fluorescence emission spectrum of the onand off state was measured with a fluorescence spectrophotometer (F-7000, Hitachi). Gel filtration chromatography was performed as described previously^16^. The molar extinction coefficient was determined by calculating the absorption values of Dreiklang and SPOON, and known protein concentration measured by an alkali denaturation method^19^. The absolute fluorescence quantum yield (φfluorescence) of Dreiklang and SPOON was measured on a Quantaurus QY (C11347, Hamamatsu Photonics), then the absorption values of Dreiklang and SPOON were adjusted to be lower than 0.05. pH titration curves were obtained as previously described^16^ and fitted by the equation described previously^20^. Photoswitching of Dreiklang and SPOON was performed by using purified proteins (10 μM in 20 mM HEPES, 150 mM NaCl buffer, pH 7.4), or the proteins expressed in *E.coli* (JM109 (DE3)) and HeLa cells. The time constant of photoswitching was measured using purified proteins embedded in 15% polyacrylamide at the final concentration and squeezed with clear cover slips to ensure the thickness of layer was ∼2 μm. We used an identical microscopy setup (inverted microscope, TE2000E) described above for imaging.. In the presence of simultaneous excitation illumination using LED light (488/25, 18 mW/cm^2^), the samples were irradiated with near-UV light (410/10, 162 mW/cm^2^) to switch the proteins from the fluorescent state to the non-fluorescent state, and UV light (360/20, 8.8 mW/cm^2^) was irradiated to switch the proteins back to the fluorescent state. The time constants (τ) for switching on and off were determined by fitting functions using single exponential growth or decay curve, respectively.

### Photoswitching by 488 nm irradiation

Photoswitching off upon 488 nm irradiation was measured by using Dreiklang or SPOON expressing *E.coli*. Measurement was done by epi-fluorescence microscopy described in single molecule localization microscopy section. Samples were irradiated by continuous 488 nm irradiation and then periodically irradiated at 100 ms with 900 ms interval controlled by a mechanical shutter system (SH05, Thorlabs). Images were taken every second. Illumination and exposure timing was controlled by LabVIEW program with a data acquisition device (NI USB-6259). Data were analyzed in Python.

### Maturation of Dreiklang or SPOON

pRSET_B_-Dreiklang or pRSET_B_-SPOON were transformed to JM109(DE3) and grown on LB plate at 37°C for overnight. After single colony picking, 10 mL LB liquid culture in 50 mL bioreactor tubes (87050, BM Equipment Co.,Ltd.) were inoculated and enclosed in a Ziploc bag with AnaeroPack (A-07, Mitsubishi Gas Chemical Company, Inc.) to remove oxygen. Then, the sample was incubated at 37°C for 42 hours. After that, the sample was chilled on ice, collected by centrifugation and then sonicated to extract protein. Followingcentrifugation, supernatants were collected, 1% penicillin-streptomycin added to prevent bacteria growth, and incubated at 37°C to induce chromophore maturation. Fluorescence was measured every 30 mins by fluorescence spectrometer (FP-750, Jusco). Data were background corrected by subtraction of the initial value and normalized by the average of last 5-time points. Data were fitted by a single exponential growth curve, using Python..

### Culture and transfection of mammalian cells

HeLa cells were cultured on 35-mm glass bottom dishes coated with collagen in Dulbecco’s modified Eagle’s medium (DMEM) supplemented with 10% fetal bovine serum (FBS). Then, HeLa cells at 50∼60% confluency were transfected with 1.0 μg plasmid DNA by using either calcium phosphate or lipofection (Lipofectamine 2000 Transfection Reagent (Life Technologies)), following manufacturer’s protocols. The culture medium was then replaced 1 hour later. The medium was washed with sterilized PBS 3 times after 8 hours and the cells were further incubated for 24-48 h in a CO2 incubator (Sanyo) at 37°C in 5% CO_2_. Imaging was performed in phenol red-free DMEM/F12 medium.

### Confocal fluorescence microscopy

Confocal imaging to confirm localization of SPOON fusion proteins was performed using a laser scanning confocal microscope (FV1000, Olympus, Japan) equipped with a multi argon-ion laser and 60× 1.35-NA oil-immersion objective lens. 515 nm light was used for the excitation of SPOON and the fluorescence signal was detected between 520-600 nm. Image processing was performed using FluoView 4.0 software (Olympus).

### Single Molecule Localization Microscopy

We used a home-built setup of objective lens epi fluorescence microscopy under inverted microscope (Ti-E, Nikon, Japan) with 100×, 1.49 NA oil-immersion objective lens (Nikon), Di01-R405/488/561/635 (Semrock) and FF01-525/45 (Semrock, USA) as dichroic mirror, and emission filter, respectively. Images were collected over an approximately 15×15 μm^2^ area with a scientific complementary metal-oxide-semiconductor (sCMOS) camera, ORCA-Flash4.0 (Hamamatsu Photonics, Japan).

### Superresolution imaging by 488 nm

HeLa cells expressing SPOON fused with vimentin were irradiated with 488 nm (1 kW/cm^2^) for 60 seconds with a frame rate of 50 Hz (20 ms). For image reconstruction, 10 to 60 seconds imaging data were used and processed with the ImageJ plugin ThunderSTORM^21^.

## Reference

1. Hell, S. W. et al. The 2015 super-resolution microscopy roadmap. J. Phys. D. Appl. Phys. 48, 443001 (2015).

2. Hell, S. W. & Wichmann, J. Breaking the diffraction resolution limit by stimulated emission: stimulated-emission depletion fluorescence microscopy. Opt. Lett. 19, 780–2 (1994).

3. Hofmann, M., Eggeling, C., Jakobs, S. & Hell, S. W. Breaking the diffraction barrier in fluorescence microscopy at low light intensities by using reversibly photoswitchable proteins. Proc. Natl. Acad. Sci. 102, 17565–17569 (2005).

4. Betzig, E. et al. Imaging intracellular fluorescent proteins at nanometer resolution. Science 313, 1642–5 (2006).

5. Rust, M. J. M. J., Bates, M. & Zhuang, X. Sub-diffraction-limit imaging by stochastic optical reconstruction microscopy (STORM). Nat. Methods 3, 793–796 (2006).

6. Gustafsson, M. G. Surpassing the lateral resolution limit by a factor of two using structured illumination microscopy. J. Microsc. 198, 82–7 (2000).

7. Tiwari, D. K. & Nagai, T. Smart fluorescent proteins: Innovation for barrier-free superresolution imaging in living cells. Dev. Growth Differ. 55, 491–507 (2013).

8. Nienhaus, K. & Nienhaus, G. U. Fluorescent proteins for live-cell imaging with super-resolution. Chem. Soc. Rev. 43, 1088–106 (2014).

9. Brakemann, T. et al. A reversibly photoswitchable GFP-like protein with fluorescence excitation decoupled from switching. Nat Biotech 29, 942–947 (2011).

10. Jensen, N. A. et al. Coordinate-targeted and coordinate-stochastic super-resolution microscopy with the reversibly switchable fluorescent protein dreiklang. ChemPhysChem 15, 756–762 (2014).

11. Hell, S. W. & Kroug, M. Ground-state-depletion fluorescence microscopy: a concept for breaking the diffracition resolution limit. Appl. Phys. B Lasers Opt. 60, 495–497 (1995).

12. Bretschneider, S., Eggeling, C. & Hell, S. W. Breaking the diffraction barrier in fluorescence microscopy by optical shelving. Phys. Rev. Lett. 98, (2007).

13. Wäldchen, S., Lehmann, J., Klein, T., van de Linde, S. & Sauer, M. Light-induced cell damage in live-cell super-resolution microscopy. Sci. Rep. 5, 15348 (2015).

14. Zhao, H., Giver, L., Shao, Z., Affholter, J. A. & Arnold, F. H. Molecular evolution by staggered extension process (StEP) in vitro recombination. Nat. Biotechnol. 16, 258–261 (1998).

15. Uno, S. et al. A spontaneously blinking fluorophore based on intramolecular spirocyclization for live-cell super-resolution imaging. Nat. Chem. (2014). doi:10.1038/nchem.2002

16. Tiwari, D. K. et al. A fast- and positively photoswitchable fluorescent protein for ultralow-laser-power RESOLFT nanoscopy. Nat. Methods 12, 515–8 (2015).

17. Suzuki, K. et al. Five color variants of bright luminescent protein for real-time multicolor bioimaging. Nat. Commun. (2016).

18. Takemoto, K. et al. SuperNova, a monomeric photosensitizing fluorescent protein for chromophore-assisted light inactivation. Sci. Rep. 3, 2629 (2013).

19. Shaner, N. C. et al. A bright monomeric green fluorescent protein derived from Branchiostoma lanceolatum. Nat. Methods 1–8 (2013). doi:10.1038/nmeth.2413

20. Shen, Y., Rosendale, M., Campbell, R. E. & Perrais, D. pHuji, a pH-sensitive red fluorescent protein for imaging of exo- and endocytosis. J. Cell Biol. 207, 419–432 (2014).

21. Ovesný M., Kř í žek, P., Borkovec, J., Švindrych, Z. & Hagen, G. M. ThunderSTORM: A comprehensive ImageJ plug-in for PALM and STORM data analysis and super-resolution imaging. Bioinformatics 30, 2389–2390 (2014).

